# Activity and abundance of nitrous oxide consuming bacteria in *Platismatia glauca* cryptogamic lichen in boreal Finnish spruce forest

**DOI:** 10.1101/2023.05.09.539975

**Authors:** Vincenzo Abagnale, Carlos Palacin-Lizarbe, Dhiraj Paul, Johanna Kerttula, Henri M.P. Siljanen

## Abstract

The boreal spruce forest soil can assimilate atmospheric N_2_O through symbiotic relationships with mycorrhizae or with bacteria, especially during spring and autumn, when aerobic microsites to soil can form. In cold soils with large field capacity (FCD), high humidity and absence of fertilisation, a balance between absorption and emission of nitrous oxide and dinitrogen was observed to be close to zero, and even to assume negative values in some cases, thus suggesting that forest soils absorb more N_2_O than they emit. Furthermore, in the presence of cryptogamic coverings of mosses and lichens, the absorption value was observed to be greater than in forests with less coverage; although the main role in N_2_O absorption is played by soil and root system. However, the role played by epiphytic organisms in N_2_O absorption in the boreal forests has not been uncovered yet. We studied, N_2_O dynamics of the lichen, *Platismatia glauca*, showing that N_2_O is consumed especially at lower incubation temperatures. The quantitative analysis with real-time PCR of nitrous oxide reductase gene fragment nosZ, showed that enzyme is present in the lichen and the gene is more transcribed under lower incubation temperature. The presented results unveil that cryptogamic covers consume nitrous oxide (with values between 0.1 and 0.4 ng N_2_O-C/g (ww)/h) at the atmospheric concentration via complete dissimilatory denitrification when nitrogen is limited.

## Introduction

The atmosphere of the shrub horizon and cryptogamic covers are consuming nitrous oxide (Machacova et al., 2017; Moyes et al., 2016); however the community of organisms that allow the reduction and assimilation of this compound is not known yet. Surely the organism in question must have access to nitrous oxide, because the reducing enzyme like NosZ/cNosZ is the only known sink for N_2_O (Pauleta et al., 2019, 2013; Ward, 2015). Another important condition depends on the enzyme, as nosZ definitely requires an anoxic environment to carry out the final denitrification step (Martens, 2005). All these characteristics must be inserted in an environment, boreal forest, with a marked seasonality, e.g. with a wide range of temperature (−40°C to +30°C), which means that the organism in question must have an active metabolism during and after these climatic conditions. In boreal forests, the fruticose and foliose lichens grow along the surfaces of the branches of the trees with most of the talus in close contact with the atmosphere. Lichens are symbiotic associations between mycobionts and photobionts. This symbiosis arises from the mutual need to maximise environmental resources usage. Nitrous oxide (N_2_O) reduction is an energetic metabolism for microbes. Usually, protoxide-reducing organisms catalyse this reaction in the absence of oxygen, therefore the creation of anaerobic pockets by the mycobiont would create a favourable microhabitat for nitrous oxide reducers (Pasqua et al., 2011).

## Material and method

### Samples collection

The collection of samples was carried out on May, 2022, in three different sites of boreal forest in Kuopio, Northern Savo, Finland: Puijo Forest (62.908328, 27.659551); Jynkänvuoren forest (62.845361, 27.694203); Kolmisoppi forest (62.873750, 27.598056) (Figure 1). *Platismatia glauca*-rich twigs were selected and collected. The samples were immediately taken to the laboratory for preparation. In addition, a 50 mL test tube of *Platismatia glauca* lichens was collected at each site; these lichens were detached from the branches on site, placed in a sterile test tube, and instantaneously frozen at -80°C. For freezing, the tubes were immersed immediately in a tank with liquid nitrogen; the samples were transported to the laboratory and stored at a constant temperature of -80°C.

**Figure 1:**
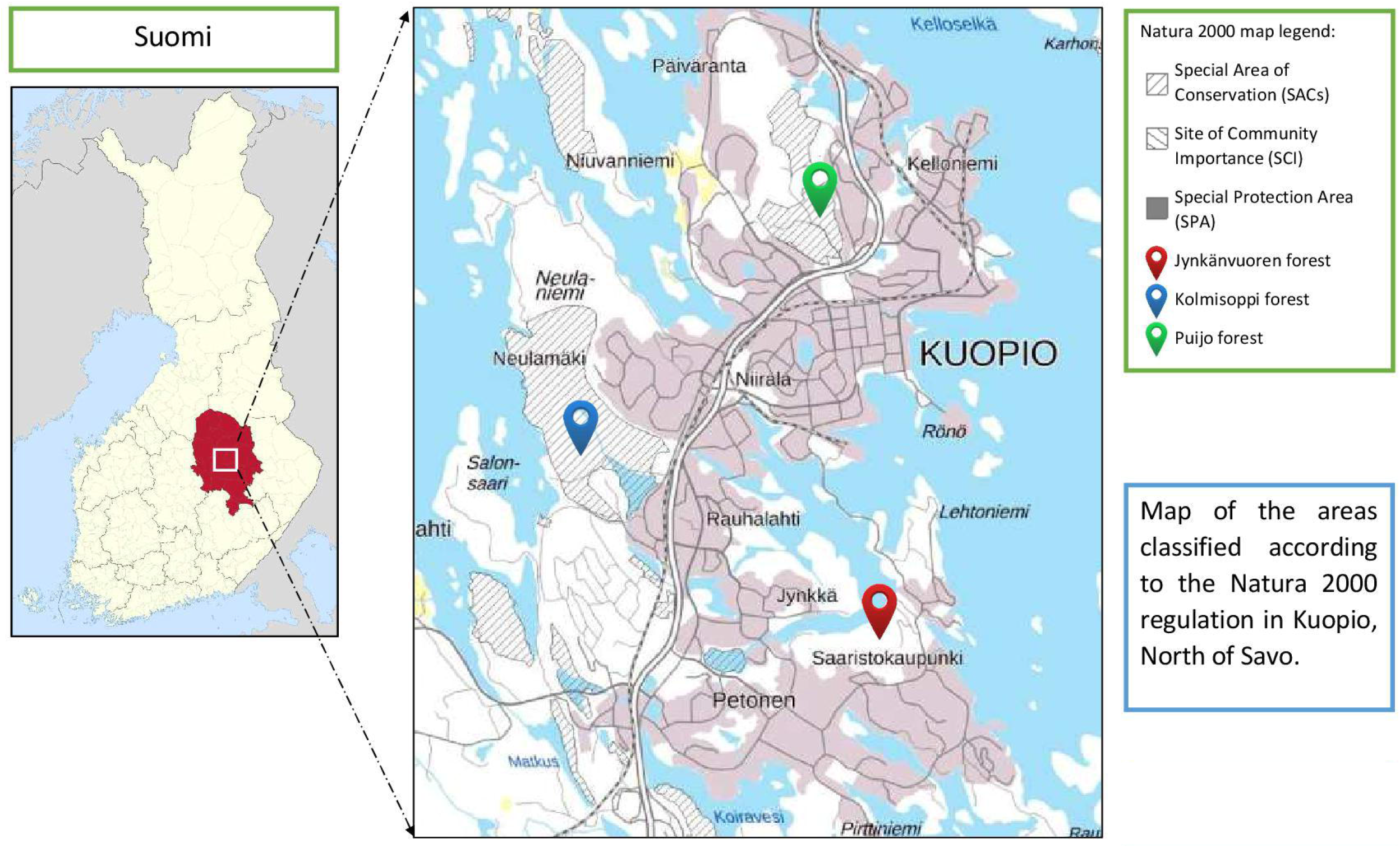
Map of sampling areas in Kuopio.

*Platismatia glauca* species samples were removed from the collected twigs with the aid of tweezers; the lichens were placed on paper to dry overnight. A bottle was prepared as in the previous test, being filled with melted snow and evacuated with the aid of helium via the evacuation line. On May 05th, 18 bottles (550 mL) were filled with about 3.01-3.06 g of lichens.

The bottles in anoxic conditions were prepared by evacuating them twice, with 1-h interval, through the evacuation line with the aid of Helium (He). The bottles in oxygenated (21% oxygen in headspace) conditions were evacuated in the same way, but then 60 mL of the air coming from the laboratory was injected in them. When necessary, sterile filtered (0.22 micrometre) and degassed melted snow was added to the samples.

The addition of N_2_O/CH_4_ gas was injected to the bottles at the beginning of the incubation with a final concentration of 20 ppm of CH_4_ and 3 ppm of Nitrous oxide. 60 mL of He were inserted into the bottles under anaerobic conditions, while additional 60 mL of air taken from the laboratory were inserted into the bottles under aerobic conditions; all bottles were incubated at 1 bar overpressure.

Gas samples were taken from the bottles by aspirating 20 mL from the headspace with a syringe and transferred to a pre-evacuated vial with a negative pressure of -1 bar. Samples were taken 1, 20, 46 and 70 hours after the start of incubation; between each sampling, the bottles were always kept in total darkness and at a constant temperature (5°C or 15°C). After completing the last sampling, the vials were analysed with gas chromatography by alternating the vials containing the headspace for each sampling time with the standard vials prepared with 20 mL of nitrous oxide at 5 ppm, methane at 15 ppm, carbon dioxide at 4000 ppm. The gas chromatograph used is an Agilent Technologies 7890B with FID1A/ECD2B/TCD3C detector. The data obtained in units of concentration were converted to ppm, and from these values the absorption curves of the various groups of samples were obtained. Data were normalised for lichen mass; in the 100% water holding capacity (WHC) samples, the data were processed in order to remove the solvation contribution of the gases in the water, so as to find the data as close as possible to the N_2_O consumption dynamics related to the lichen only.

### DNA and RNA extraction

After having carried out the headspace sampling, the samples were immediately frozen in liquid nitrogen to block any metabolic process. Each bottle was opened and emptied of its contents into a sterile mortar, where the lichens were pulverised with a pestle while maintaining a low temperature thanks to the continuous mixing with liquid nitrogen. The lichen powder was then transferred into pre-labelled 20mL test tubes by using sterile spatulas, and frozen in a cold room at -80°C. The same procedure was used for the three samples frozen directly in the field.

DNA was extracted by a phenol/chloroform/isoamyl alcohol double cycle procedure and purified by RNase; RNA extraction was performed with a kit called RNEasy Plant Mini Kit produced by Qiagen; in particular the protocol related to plant tissues and fungi was used (QIAGEN, 2018). The extracted RNA was purified by using an RNase-free DNase solution (Thermo scientific DNase I, RNase-free) in order to completely eliminate the DNA from the extract without digesting the RNA; the latter was PCR-assayed with nosZ-related oligos to detect possible DNA that was not hydrolysed by DNase, then, as no DNA amplifications were found, the RNA could be retro transcribed into cDNA.

### cDNA synthesis

The purified RNA was used for a reverse-transcription step in order to produce complementary DNA (cDNA)ì. The cDNA was produced with MuLV H-minus transcriptase (Maxima MuLV H minus, Fermentas, Lithuania) and random hexamers. First the random hexamers (200 ng/µL) were added on the top of the RNA (7 µL volume of DNA-degraded RNA). The random hexamers were induced to anneal with RNA at +65°C, for 5 minutes. Then, the reverse transcriptase buffer was added, Ribolock RNase inhibitor and 1µL reverse transcriptase H minus. The reverse transcriptases were incubated at +25°C for 10 minutes and then at +50°C for 30 minutes.

### q-PCR and RT-qPCR

The DNA and cDNA extracts of the dry samples were selected to be amplified with specific primers, namely those targeting the *nosZ* gene of Clade 1 and Clade 2 (Tab. 1).

The primers were selected in order to reveal how abundant Clade 1 and Clade 2 were in the samples (Henry et al., 2006; Jones et al., 2013). The sequencing of the samples was done with primers having Illumina sequencing adapters; these adapters are used in sequencing service laboratories to index the samples for Illumina sequencing. The PCR reaction and cycling conditions selected are shown in Tables 2 and 3.

**Table 1:**
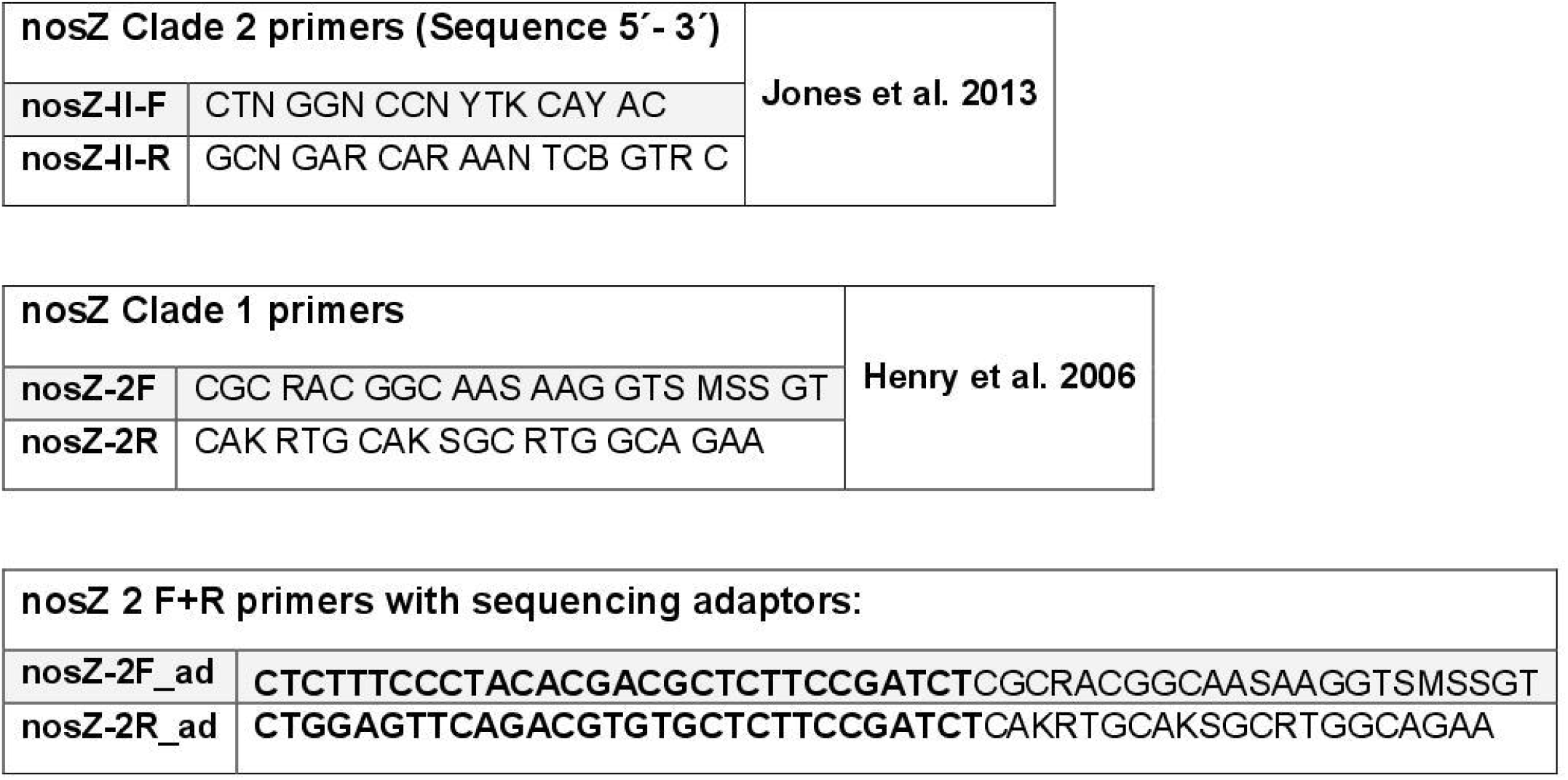
List of used primers.

**Table 2:**
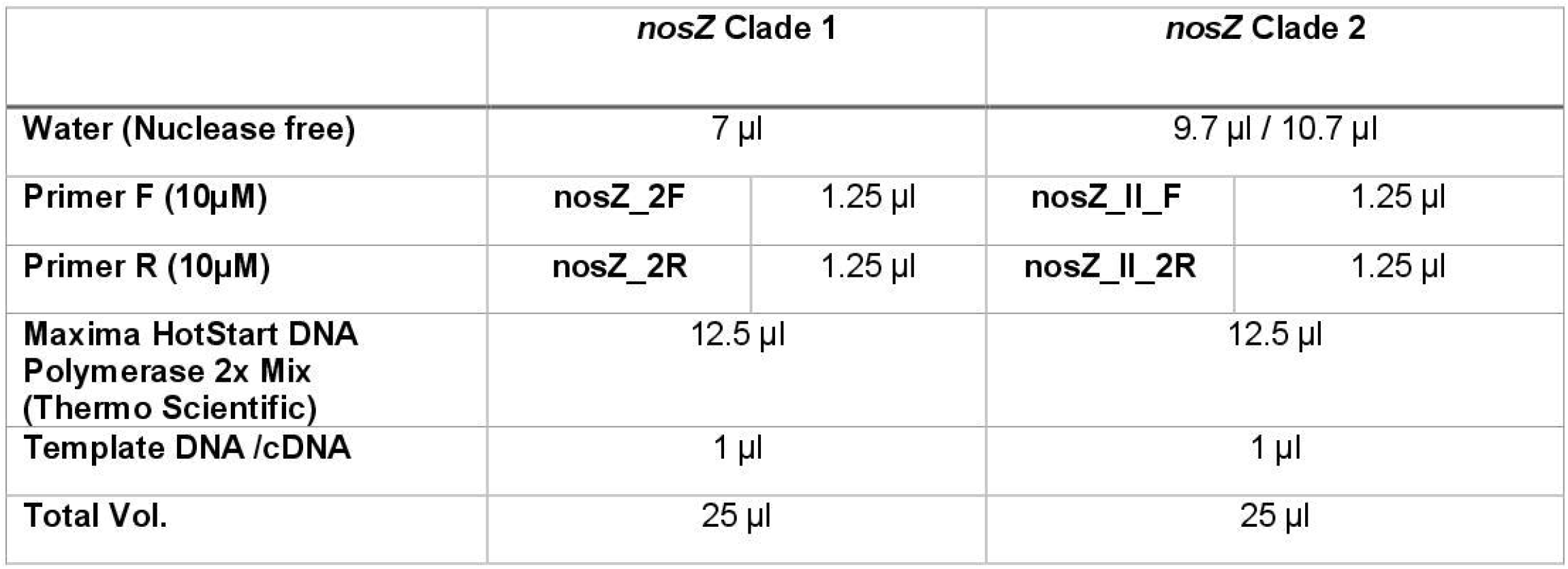
PCR Mixtures

| **Table 2: PCR Mixtures** | ***nosZ* Clade 1** |  | ***nosZ* Clade 2** |  |
| --- | --- | --- | --- | --- |
| **Water (Nuclease free)** | 7 μl | | 9.7 μl / 10.7 μl | |
| **Primer F (10μM)** | ***nosZ_2F*** | 1.25 μl | ***nosZ_II_F*** | 1.25 μl |
| **Primer R (10μM)** | ***nosZ_2R*** | 1.25 μl | ***nosZ_II_2R*** | 1.25 μl |
| **Maxima HotStart DNA Polymerase 2x Mix (Thermo Scientific)** | 12.5 μl | | 12.5 μl | |
| **Template DNA /cDNA** | 1 μl | | 1 μl | |
| **Total Vol.** | 25 μl | | 25 μl | |

**Table 3:**
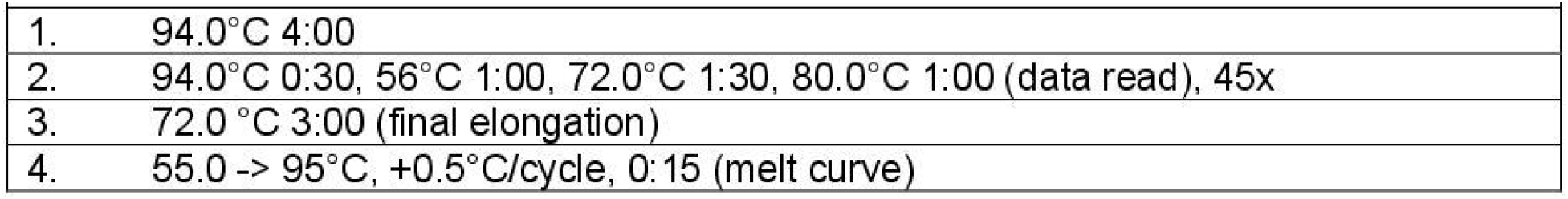
PCR cycling conditions

Quantitative PCR was done with a Bio-Rad MyCycler device. Quantification was done with duplicate reactions per sample. For sequencing, three PCR products were pooled per sample. The PCR was done with amplification shorter than 35 cycles, in order to avoid the production of chimeric sequences. The products for sequencing were purified with a PCR product purification kit (Roche, High Pure PCR Product-kit), according to the manufacturer’s protocol, after having verified the specificity of all PCR products by gel electrophoresis; concentration was measured with NanoDrop spectrophotometer.

## Results

### GC analysis

The data obtained from gas chromatographic analysis (Figure 2) show us that in the presence of oxygen *Platismatia glauca* tends to consume nitrous oxide, whereas in the absence of oxygen, consumption and emission fluxes tend to be equal (0.006 ± 0.010 ng N_2_O-C/g (ww)/h in dry samples and 0.051 ± 0.034 ng N_2_O-C/g (ww)/h in hydrated samples). The highest consumption of nitrous oxide was identified in hydrated samples at 5°C in the presence of oxygen with consumption fluxes of -0.407 ± 0.071 ng N_2_O-C/g (ww)/h; increasing the incubation temperature, as in samples at 15°C, the consumption flux is lower (−0.110 ± 0.089 ng N_2_O-C/g (ww)/h in dry samples and -0.104 ± 0.064 ng N_2_O-C/g (ww)/h in hydrated samples) while inoculating at 5°C with dehydrated lichens also results in a lower consumption flux of -0.104 ± 0.054 ng N_2_O-C/g (ww)/h. The variations in the concentration of N_2_O in the headspace of the samples depend on the fact that the system changes punctually going to consume or produce nitrous oxide. In fact, we found that at high concentrations of nitrous oxide subsequently the system tends to consume while when nitrous oxide concentrations are low, the system tends to emit; It was found that the system in 550 ml bottles under aerobic conditions tends to volume between 230-240 ng N_2_O-C/g, while the same system under anoxic conditions tends to volume between 130-150 ng N_2_O-C/g. These processes are influenced by N_2_O concentration itself, thus leading us to deduce that the consumption activity is continuously activated and deactivated depending on the most available substrate. In fact, the variation of metabolic activities is the result of the presence/absence of various substrates which can be organised, reduced, and oxidised in the environment; in particular, carbon-based organic and inorganic molecules allow or prevent the activation of various pathways based on the concentration of the substrates.

**Figure 2:**
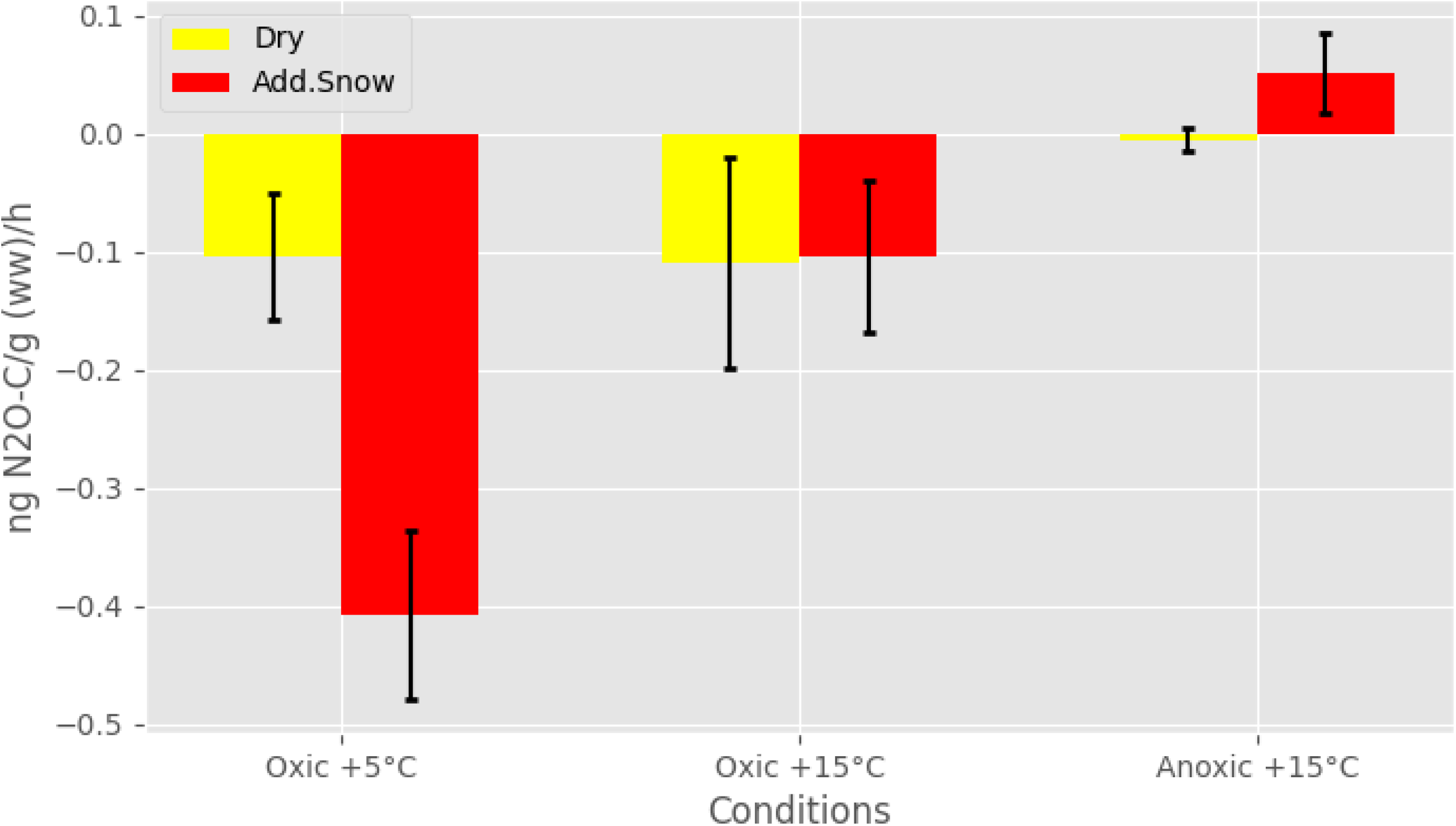
Platismatia glauca N2O flux variables in air condition and temperature.

### qPCR and RT-qPCR results

All the samples demonstrated the presence of *nosZ-*carrying microbes in the microbiome associated with *Platismatia glauca* (Fig.3). In particular 4.04 ±3.30 E+05 copies of the gene were counted per gram of lichen *in situ*, while the samples inoculated in the laboratory demonstrated a greater number of copies of the gene corresponding to:

**Figure 3:**
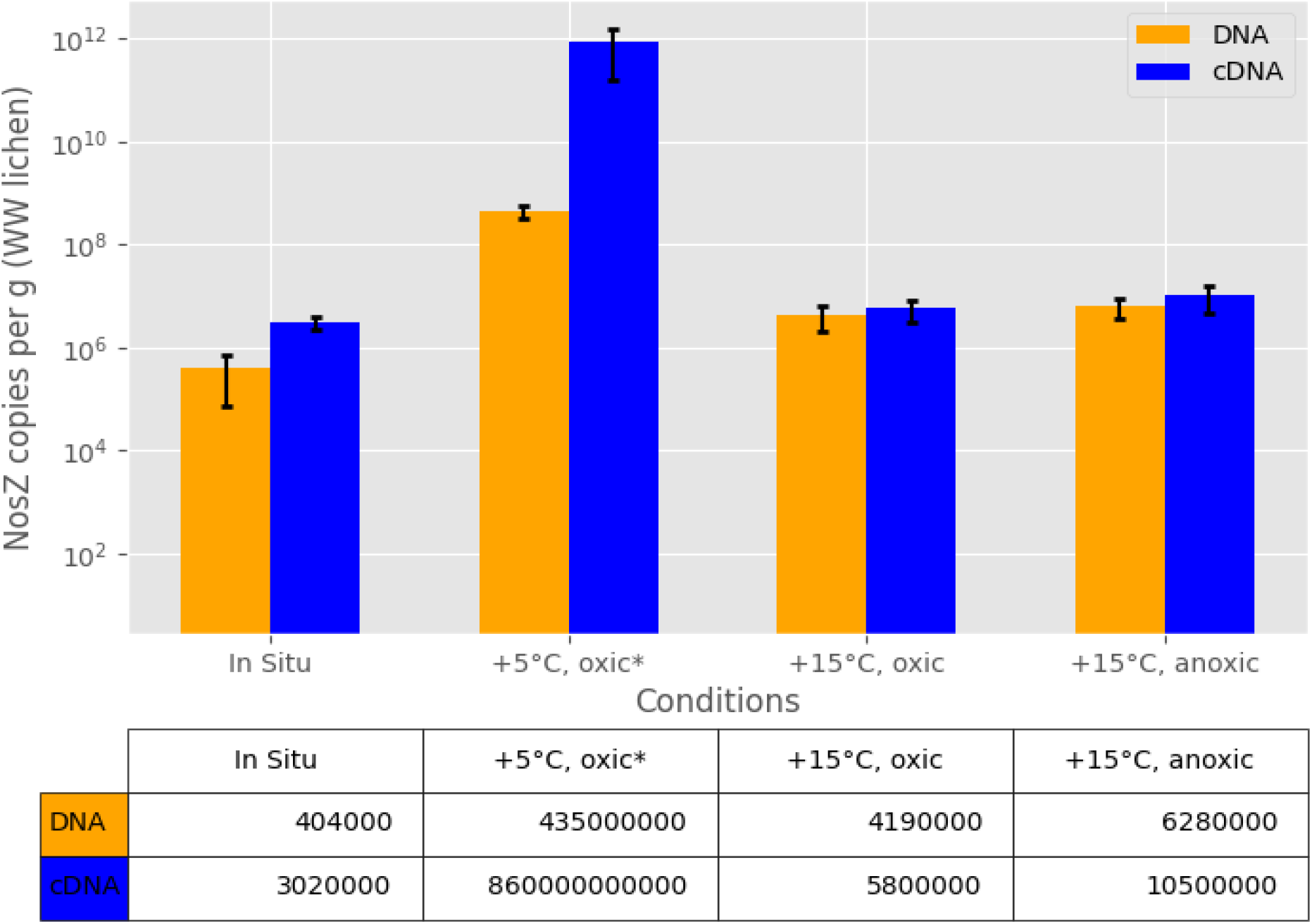
Abundance of nosZ gene copies (in orange) and transcription (in blue) in *Platismatia glauca*. Statistical pairwise comparison to the *in-situ* samples is shown with asterisk (P < 0.05) [MOU1]. The mean and standard deviation of three biological replicates (n = 3) is shown.

- 4.19 ± 2.21 E+06 copies of *nosZ* clade I per gram of lichen in samples treated at 15°C in an aerobic environment (about one order of magnitude greater than *in situ* measurement);
- 6.28 E+06 ±2.78 E+06 of *nosZ* clade I copies per gram of lichen in samples treated anaerobically at 15°C (about one order of magnitude greater than *in situ* measurement),
- 4.35 E+08 ±1.15 E+08 copies of *nosZ* clade I per gram of lichen in samples treated at 5°C in an aerobic environment (about three orders of magnitude greater than the measurement). The gene identity was confirmed by sequencing the nosZ gene, with amplifying the samples with the adapter primers of Illumina Miseq sequencing platform and library preparation and sequencing was done in Azenta company (Genewiz, Azenta, UK).

The data therefore indicate that total darkness favours N2O-reducing microbes, by enhancing their abundance and the expression of the *nosZ* gene. As regards to the data related to *in situ* expression, *e*.*g*. mRNA values, only the Puijo site provided valid indications on the expression of *nosZ* (4.04 ± 3.3 E+05 copies per gram of lichen), indicating almost an order of magnitude more (3.02 ± 0.82 E+06 copies per gram of lichen) than the DNA copies. With regard to samples at different incubation temperatures, treatment at the lowest temperature (5°C vs. 15°C) results in higher nosZ gene abundance and expression (8.60 ±7.02 E+11 mRNA copies per gram of lichen vs. 4.35 ±1.15 E+08 DNA copies per gram of lichen), suggesting that nosZ-carrying microbes have a low temperature optimum. As far as clade II of *nosZ* is concerned, the qPCR gave completely negative results both from the point of view of gene expression and as regards the presence of the gene itself.

## Discussion

Returning to the objectives of this study, Platismatia glauca under any temperature and hydration condition tested performs nitrous oxide consumption dynamics. This phenomenon can be explained by the need of Cephalodium cyanobacteria (Brodo et al., 2002) to release cytochromes from electron accumulation due to forced metabolism in anaerobiosis. Nitrous oxide can certainly be a valuable electron acceptor, not really because of its electron acceptor capacity, but because of the ease of availability in anoxic systems given incomplete dissimilative denitrification processes, said formation of anaerobic pockets in the lichen by the mycobiont promotes complete denitrification phenomena from nitrate to molecular nitrogen.

Lichens exploit an evolutionary advantage, namely the mycobiont-photobiont symbiosis, which allows the storage of water and nutrients, the capture of simple substances directly from the atmosphere in the form of gases or through precipitation as solutes, and their processing through a large pool of metabolic pathways such as photosynthesis, denitrification, methanotrophy, etc. Given the large number of reactions, we sought to understand what factors really influence the activation of the nosZ pathway. Reading previous studies on Finnish forests, peak nitrous oxide consumption has generally been recorded during the snow season, particularly between October-November and February-March; during this period there are few and low light hours. These results are understandable, given the transition from photosynthetic to chemosynthetic metabolism. Although with lower intensity, denitrification of nitrous oxide through the N_2_OR pathway is also present in brightness, as demonstrated by test in situ samples.

A key factor in the expression of *nosZ* and thus in the dynamics of denitrification is the temperature. Indeed, experiments have shown that a decrease in temperature corresponds to increased activation of the pathway; however, the motivation for this behaviour is still unclear.

A regulatory mechanism of the lichen itself could be hypothesised, as near-zero temperatures reduce the availability of nitrate substrates; therefore, the lichen could increase the activation of Nitrous Oxide Reductase to use nitrous oxide, which is more available and in gaseous form, as an electron acceptor. Temperature also changes the solubility of gases in water; at lower temperatures, the solubility of gases is greater than at higher temperatures (Palacin-Lizarbe et al., 2018).

One factor that initially caused confusion is the degree of hydration of the lichen. Untreated water from snow is indeed rich in nitrate and ammonia, so it could trigger the mechanisms of ammonia nitrification and nitrate denitrification. The combination of these pathways may increase nitrous oxide emission fluxes, which, however, were partially consumed by the N_2_OR complex (Pauleta et al., 2019). To verify this possibility, in the test, water was filtered with a 0.22 μm microfilter and sequentially evacuated with helium. Filtration removed micro and nanoparticles of wood or other organic matter, while outgassing removed dissolved gases by varying atmospheric pressure values. The end result, in lichens hydrated by this technique, was a higher consumption of nitrous oxide, justified by the presence of nitrate in insufficient quantities to overcome the electronic requirements of the electron transport chain.

With these statements and hypotheses, it can be understood that lichen hydration contributes as a temperature cofactor to the dynamics of N_2_O consumption, mainly favouring the exchange of substances in the periplasm within the lichen, logically if the water absorbed by the mycobiont is rich in nitrates and ammonia, the nitrification and incomplete denitrification pathways will be favoured. With regard to nitrous oxide dynamics in Savo’s forests in summer, the conditions are as follows:

- temperatures close to 20°C
- high levels of light radiation
- decomposition events of organic matter
- utilisation by eukaryotic and prokaryotic organisms of previously mineralised substances,
- nitrification and anammox processes with increased nitrate concentrations,

result in the emission of nitrous oxide and carbon dioxide. During this period, lichens have optimal light conditions, so they can photosynthesise organic substances through the algal layer and utilise energetically convenient pathways for electron transport chains.

When temperatures fall near or below 0°C, there is little light and not much free organic and inorganic matter, because it is immobilised in the soil. Cryptogamic covers and some denitrifying bacteria present on conifer leaves can survive these conditions by producing organic molecules that exploit nitrous oxide as an electron acceptor with an energy yield of ATP, to be used in anaerobic processes such as the Calvin cycle (Dehò and Galli, 2012). Symbiotic links are possible between lichen cells and microbial cells, which can produce energy for anaerobic gluconeogenesis. Although our data hint at this direction, they cannot confirm it.

As regards the experiments in anaerobic conditions, we have ascertained that it does not cause the cryptobiosis of the lichen, as nitrous oxide emission dynamics were observed. In addition, despite the need for anaerobic pockets for denitrification, *Platismatia glauca* does not increase the expression of *nosZ* gene in the total absence of oxygen, but in aerobic conditions at high hydration and low temperatures. It can therefore be hypothesised that a greater expression of *nosZ* is attributable to aerobic pathways that need fast electron acceptors: when the periplasm is saturated with nitrous oxide, it becomes the most available substrate, and Cyanobacteria increase the expression of *nosZ* up to when nitrous oxide returns to non-saturating levels. In fact, a first phase of high nitrous oxide consumption was observed in the first hours of aerobic incubation of *Platismatia glauca*, followed by a continuous phase of equilibrium between gas consumption and emissions, seen as a sort of research of stability by the lichen.

Considering the results, in general, it can be confirmed that the lichens of the *Platismatia glauca* species and the associated microbiota allow the reduction of nitrous oxide into molecular nitrogen in aerobic conditions, confirming that the uptake can take place in natural condition thanks to enzymes encoded by the gene *nosZ* of clade I organism attributable to imperfect denitrification events (Graf et al., 2016). Nonetheless, nitrous oxide is also absorbed in anaerobic conditions, with the difference that the balance between gas emissions and consumption favours emissions. In fact, the gas chromatographic values identify either the emission or balanced dynamics between the production and absorption of nitrous oxide. As regards the number of copies of DNA and the amount of expression in RNA, in anaerobic conditions the rate of in RNA and translation in enzymes is similar to that observed in aerobic conditions; however, the number of copies of sequences is lower by two orders of magnitude.

The variables favourable to the absorption of N_2_O can therefore be identified in:

1. the low temperatures, especially at values below 5°C;
2. the absence of light sources;
3. the conditions of high relative humidity (the values tested were close to 100% saturation);
4. the presence of oxygen.

All this makes us understand that N_2_O consumption generally develops aerobically, that the limiting factor is the presence of oxygen, that the accelerating factor is the low temperature (T ⪝ 5°C), that water is a cofactor of temperature; these factors and cofactors, on the basis of given values, activate/deactivate genes encoding the genes involved in N_2_O-related metabolic pathways present in lichens and microbial communities connected to them.

Ultimately, the data indicate that the *nosZ* gene fragment of clade I is present in the microbiome associated with *Platismatia glauca* at basal levels and that its abundance and expressivity can increase with the onset of low temperatures below 5°C. These results are in agreement with the data related to the nitrogen cycle in boreal forests, in which the drop in temperatures induces the loss of nitrogen and carbon due to their immobilisation in the soil. Therefore, organisms possessing secondary metabolic pathways can activate them as a survival mechanism.

Instead, a hypothetical explanation regarding the positive finding in the genome but negative in the transcriptome of *nosZ* in the *in situ* samples examined taken from Jynkänvuoren forest and Kolmisoppi forest can be attributed to the photosynthetic possibility of the lichens themselves. In Puijo the luminosity is scarce even in May given the greater tree cover, while in Kolmisoppi and Jynkänvuoren the covers are much more open given the young age of the trees. A lower availability of light translates into a lowering of the energy production derived from photosynthesis; through the Cyanobacteria of the Cephalodium, the lichen could exploit this anaerobic pocket to carry out the complete dissimilatory denitrification up to the release of molecular nitrogen, thus producing membrane proton gradient useful for the formation of ATP without affecting the aerobic pathways of the lichen itself.

It can be hypothesised that in the boreal forests, during the dark season, with temperatures close to or below 0°C, photosynthesis is blocked due to the lack of light, and fermentation-type metabolism is blocked due to the lack of low weight organic substrates molecular; this block therefore favours the energy pathways that use simple molecules such as ammonia and nitrate. Ammonia is oxidised by ammonia monooxygenase (AMO) and hydroxylamine oxidoreductase (HAO) to nitrite, which will be denitrified into nitric oxide or nitrified into nitrate (Dehò and Galli, 2012; Sayavedra-Soto et al., 1996). Denitrification reduces nitrate to NO by via the quinones, thus allowing the acidification of the periplasmic space and therefore the energy yield in ATP via the H+ proton pump; however, the electron transport chain can become clogged on the cytochrome *bc* complex due to the lack of acceptors of electrons and the accumulation of substrates such as NO_2_^-^, NO and N_2_O (Lee et al., 2022). To remedy the shortcoming, the cell transcribes and translates the *nosZ* gene which can accept two electrons for each nitrous oxide molecule, reducing it to N_2_ with the release of water.

## Conclusion

Cryptogamic covers have a role in the dynamics of nitrous oxide, as all four lichen species studied show the capability of consuming N_2_O. The epiphytic lichens present in the boreal horizons present complete dissimilatory denitrification pathways. The abundance of gene and transcript copies of the *nosZ* fragment at basal levels was found in all *Platismatia glauca* samples; an enrichment in gene copies and expression was observed in the samples at 5°C in oxic and dark conditions.

