## Supplementary material for "Activity and abundance of nitrous oxide consuming bacteria in *Platismatia glauca* cryptogamic lichen in boreal Finnish spruce forest": pdf_supplemental

All this makes us understand that $N_2O$ consumption generally develops aerobically, that the limiting factor is the presence of oxygen, that the accelerating factor is the low temperature ($T \lesssim 5°C$), that water is a cofactor of temperature; these factors and cofactors, on the basis of given values, activate/deactivate genes encoding the genes involved in $N_2O$-related metabolic pathways present in lichens and microbial communities connected to them.

Table 1: List of used primers.

| nosZ Clade 2 primers (Sequence 5´- 3´) | | |
|---|---|---|
| **nosZ-II-F** | CTN GGN CCN YTK CAY AC | **Jones et al. 2013** |
| **nosZ-II-R** | GCN GAR CAR AAN TCB GTR C | |

| nosZ Clade 1 primers | | |
|---|---|---|
| **nosZ-2F** | CGC RAC GGC AAS AAG GTS MSS GT | **Henry et al. 2006** |
| **nosZ-2R** | CAK RTG CAK SGC RTG GCA GAA | |

| nosZ 2 F+R primers with sequencing adaptors: | |
|---|---|
| **nosZ-2F_ad** | **CTCTTTCCCTACACGACGCTCTTCCGATCT**CGCRACGGCAASAAGGTSMSSGT |
| **nosZ-2R_ad** | **CTGGAGTTCAGACGTGTGCTCTTCCGATCT**CAKRTGCAKSGCRTGGCAGAA |

Clade 1 primers are
nosZ-2F
nosZ-2R, Henry et al., 2006.

| Table 2: PCR Mixtures | *nosZ* Clade 1 | | *nosZ* Clade 2 | |
|---|---|---|---|---|
| **Water (Nuclease free)** | 7 µl | | 9.7 µl / 10.7 µl | |
| **Primer F (10µM)** | **nosZ_2F** | 1.25 µl | **nosZ_II_F** | 1.25 µl |
| **Primer R (10µM)** | **nosZ_2R** | 1.25 µl | **nosZ_II_2R** | 1.25 µl |
| **Maxima HotStart DNA Polymerase 2x Mix (Thermo Scientific)** | 12.5 µl | | 12.5 µl | |
| **Template DNA /cDNA** | 1 µl | | 1 µl | |
| **Total Vol.** | 25 µl | | 25 µl | |

| **Table 3: PCR cycling conditions** | |
|---|---|
| 1. | 94.0°C 4:00 |
| 2. | 94.0°C 0:30, 56°C 1:00, 72.0°C 1:30, 80.0°C 1:00 (data read), 45x |
| 3. | 72.0 °C 3:00 (final elongation) |
| 4. | 55.0 -> 95°C, +0.5°C/cycle, 0:15 (melt curve) |

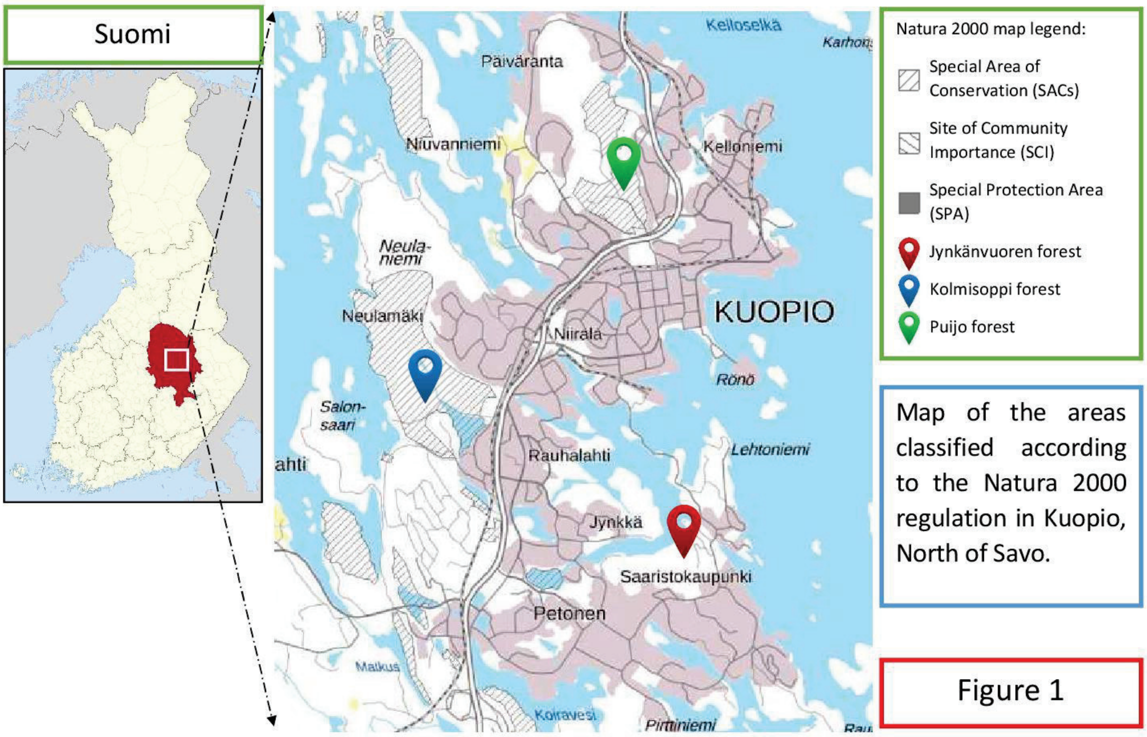


Figure 1: Map of sampling areas in Kuopio.

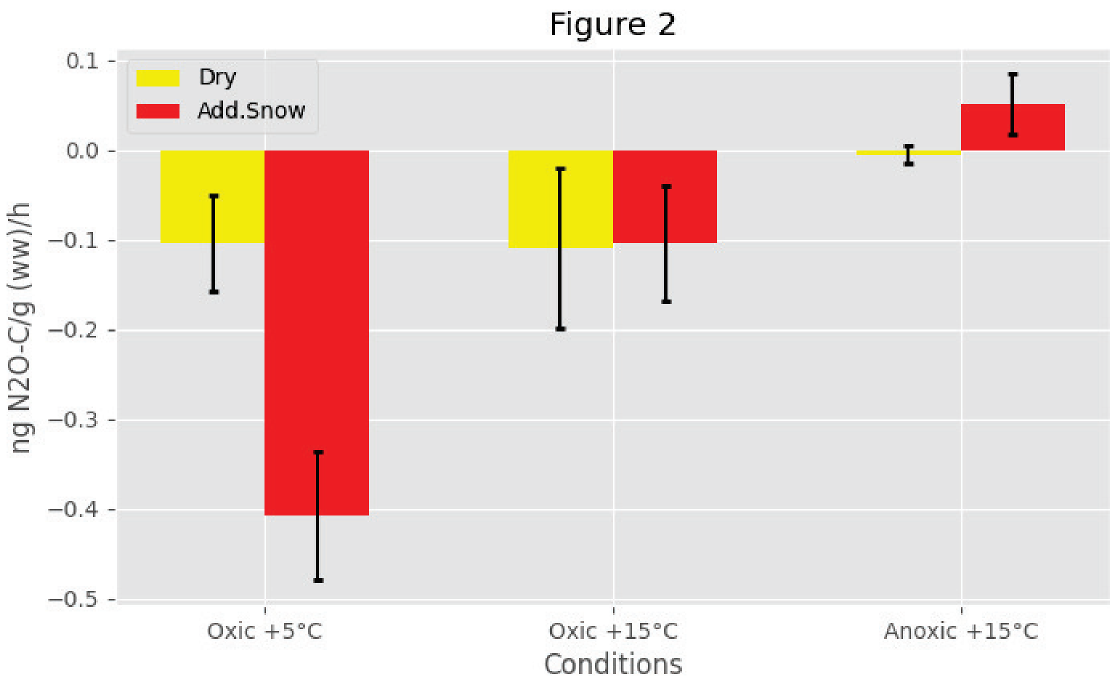


Figure 2: *Platismatia glauca* $N_2O$ flux variables in air condition and temperature.

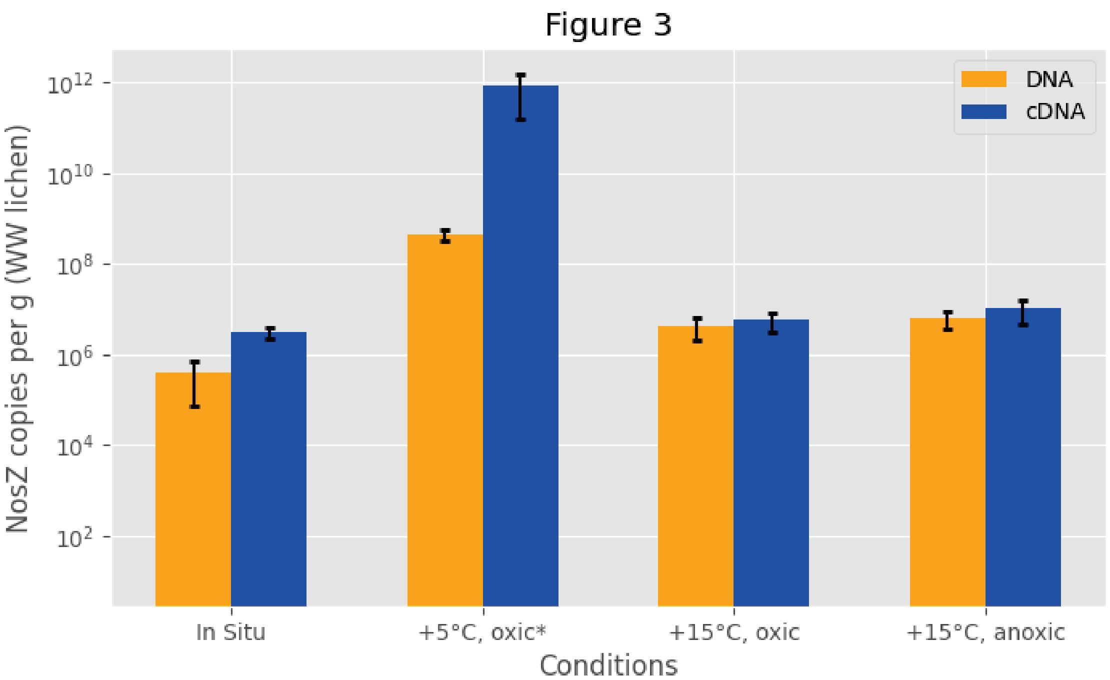


| | In Situ | +5°C, oxic* | +15°C, oxic | +15°C, anoxic |
|---|---|---|---|---|
| DNA | 404000 | 435000000 | 4190000 | 6280000 |
| cDNA | 3020000 | 860000000000 | 5800000 | 10500000 |

Figure 3: Abundance of nosZ gene copies (in orange) and transcription (in blue) in *Platismatia glauca*. Statistical pairwise comparison to the *in-situ* samples is shown with asterisk ($P < 0.05$) [MOU1]. The mean and standard deviation of three biological replicates ($n = 3$) is shown.
